# A spatial analysis of Common Pheasant (*Phasianus colchicus*) abundance with reference to Protected Area coverage in England

**DOI:** 10.64898/2026.05.06.722883

**Authors:** Joe A. Wilde, Luke Ozsanlav-Harris, Joah R. Madden

**Affiliations:** Center for Research in Animal Behaviour, Psychology, University of Exeter, Exeter, EX4 4QG, UK; Biomathematics and Statistics Scotland, The James Hutton Institute, Aberdeen, AB15 8QH, Scotland, UK; University of Exeter, Penryn Campus, Penryn, Cornwall, TR10 9FE, UK

**Keywords:** Pheasants, Hunting, Protected Areas, Ecological effects, Dispersal, Survival

## Abstract

The release of tens of millions of common pheasants (*Phasianus colchicus*) across the UK for shooting may pose an ecological risk to native species and sensitive habitats, particularly if the birds move into protected areas (PAs) such as Special Areas of Conservation (SAC), Special Protection Areas (SPA), and Sites of Special Scientific Interest (SSSI). The extent of this ecological risk depends on the abundance of pheasants in these sensitive sites, especially if they are attracted there after the shooting season when game management efforts to retain the birds cease. We used relative pheasant abundance measures derived from British Trust for Ornithology bird atlas data from 3793 2km tetrads across four English counties (Berkshire, Cornwall, Devon, and Hertfordshire) to determine if pheasants preferentially disperse into or reside in areas with greater PA coverage. We analysed relative abundance in both the winter shooting season and the breeding season using a Bayesian occupancy-abundance model, controlling for habitat type and diversity. Our results showed a strong influence of habitat on pheasant abundance, consistent with known habitat preferences. However, we found no evidence of a relationship between relative pheasant abundance and the proportion of ecologically relevant PA coverage in a tetrad. This lack of a relationship was consistent across all four counties and across both the winter and breeding seasons. Our finding suggests that common pheasants do not preferentially disperse into or reside in protected areas compared to surrounding, unprotected land, suggesting that the ecological impacts caused by released pheasants are no more likely to occur in protected areas than in non-protected areas.

## INTRODUCTION

Tens of millions of common pheasants (*Phasianus colchicus*) are released across the UK in late summer/early autumn for shooting (Madden, 2021). Recent estimates of release numbers range from 31.5 million pheasants (range 29.8–33.7 million; Madden, 2021) or 47 million pheasants (39-57 million 95% confidence interval; Aebischer, 2019). These releases occur across lowland United Kingdom from an estimated 7000-9000 sites (Madden, 2021) and it is estimated that 60% of the country is influenced by game management (PACEC 2014) and involves 28% of English woodland (Gilbert, 2007).

Once released, pheasants can move away from release sites (Hill and Ridley, 1987) but game management aims to keep the majority of pheasants within areas where shooting will take place (Hill, 1988). Although management actions that accompany the release of these birds, such as habitat creation or maintenance, supplementary feeding, and legal pest and predator control, may provide benefits to other non-game species, the direct actions of the released birds themselves are likely to be ecologically damaging, especially if best practice advice is not followed (Madden & Sage, 2020; Mason et al., 2020; Sage et al., 2020; Madden, 2023). There is evidence to suggest that released birds can increase soil and water nutrient levels, directly damage flora, predate on local species, compete for resources with non-game species, provide a source or harbour for pathogens and parasites that may infect other species (Madden & Sage, 2020; Mason et al., 2020; Sage et al., 2020; Madden, 2023). These negative effects are likely to be stronger at higher densities of released birds (e.g. Sage et al., 2005a; Gortazar et al., 2006; Pressland, 2009; Neumann et al., 2015; Porteus, 2015; Capstick et al., 2019; but see Davey, 2008). Changes in soil chemistry and plant species richness can persist at woodland release sites for around a decade after releases cease, but this may be longer if the densities of released birds are >1000 birds/ha (Capstick et al., 2019).

The direct negative effects of released gamebirds may be of particular concern if they occur in habitats that contain ecologically important features that are susceptible to damage from gamebirds. Such habitats containing ecologically important features are commonly legally protected. In England, some of the most important sites include Special Areas of Conservation (SAC, designated under the Habitats Directive) and Special Protection Areas (SPA, designated under the Birds Directive). Collectively, these form part of the Natura 2000 network that spans Europe. The UK government is under obligation to prevent deterioration of habitats in Natura 2000 sites. A third set of sites, Sites of Special Scientific Interest (SSSI), may also be subject to direct negative effects if they contain designated features susceptible to gamebird activity. Releases of gamebirds within or close to these areas are regulated (https://www.gov.uk/guidance/gamebirds-licences-to-release-them). Releases inside, or within a buffer zone of 500 metres, of Natura 2000 sites currently require a licence. This may be a general licence (GL43 for SACs, GL45 for SPAs covering pheasants and partridges) if a series of conditions relating to release densities (<700 birds/ha of release pen/area within the site or <1000 birds/ha of release pen/area within the buffer zone), release site habitats, feeding sites and reporting requirements can be met. If these conditions cannot be met, then an individual licence must be applied for (DEFRA, 2025). Releases and aspects of their subsequent management in SSSIs may need consent from Natural England (DEFRA, 2024). There is currently sparse evidence of the effects from gamebird releases specific to species and habitats that occur within protected areas (Rothero, 2006; Bosanquet, 2019; Hand, 2020). Stakeholders (including nature reserve staff, farmers and gamekeepers) hold a variety of perceptions and opinions on what the effects of release and management might be on protected areas. There is broad agreement that there are benefits of hedgerow creation and management accompanied by negative effects arising from increased generalist predator populations (Minter et al. 2024). However, there is little understanding of what the abundance of released gamebirds might be in protected areas and whether gamebirds released outside protected areas move into them.

SPAs, SACs, and SSSIs (hereafter referred to as protected areas, PAs) may be attractive to dispersing pheasants as they may provide a refuge from shooting activities or other forms of human disturbance or contain suitable wintering/breeding habitats, such that they actively seek out PAs and disperse into them. Additionally, released birds may simply diffuse from release sites and passively arrive in PAs when game management around their release areas ceases. Because birds are rarely released in these PAs, birds within them are likely due to immigration, with the potential to produce densities causing ecologically damage. It is therefore important to determine whether released pheasants preferentially disperse into protected areas to determine the risk of ecological change in these areas. While there is previous research focusing on changes in pheasant abundance in other countries (McGowan *et al*., 1999; Coates *et al*., 2017; Barg *et al*., 2025), and research focusing on the impact released gamebirds have on habitat in the UK (Sage *et al*., 2020; Neumann *et al*., 2015), we are unaware of previous research focusing on changes in pheasant abundance in relation to protected areas in the UK.

We assessed whether pheasants are moving into protected areas by examining whether there were shifts in the relative abundance of pheasants from areas where they were released and retained by game management (in late summer to late winter) towards protected areas in the breeding season (in late spring/early summer when shooting and game management has predominantly ceased). If protected areas are attractive to released pheasants, and pheasants therefore preferentially reside in them, then we might expect to find large-scale movements of pheasants from areas where they were released (and therefore reside during late summer/early winter) into protected areas, especially when game management has declined or ended in the spring following the shooting season, and activities limiting dispersal have ceased. Therefore, we are interested in changes in the abundance of pheasants (relative to the county-level local population at a given time) in relation to the amount of PA that an area contains. Investigating relative pheasant abundance allows us to account for changes in population size due to mortality and releases. If pheasants are dispersing towards protected areas, then we would expect a positive relationship between protected area coverage and relative pheasant abundance, or a positive change in that relationship between seasons. A positive relationship between protected area coverage and relative abundance in both winter and the breeding season would suggest protected areas are consistently attractive to pheasants despite efforts from game management to limit pheasant dispersal. However, this positive relationship may only be present, or be especially pronounced, during the breeding season (i.e. there may be an interaction between protected area coverage and time period), when efforts by game management to limit pheasant dispersal have ended. Alternatively, there may be no relationship or change in relationship over time, indicating pheasants do not preferentially disperse to or reside in protected areas.

## METHODS

### i) UK bird atlas data

During 2007-2011, the British Trust for Ornithology (BTO) organised volunteer surveys of all UK birds at a 2km x 2km tetrad level. These occurred during the winter (November to February) and the breeding season (April to July; Balmer et al., 2013). Fieldwork for the survey, carried out in four winters (2007/08–2010/11) and four breeding seasons (2008–11), consisted of a pair of standardized visits to each tetrad in each of the eight survey periods (for more details on the methodology see Balmer *et al*., 2013; Gillings *et al*., 2019). In some areas, there was complete coverage, with all 2km^2^ tetrads being surveyed. Such complete data was available from four counties (Figure 1) who published their UK bird atlases: Berkshire (http://berksoc.org.uk/county-atlas/, Clews 2013), Cornwall (https://atlas.cbwps.org.uk/), Devon (Beavan & Lock 2016) and Hertfordshire (http://www.hertsatlas.org.uk/, Smith *et al*. 2015). For each of these atlases, the maximum recorded abundance of birds in 2km x 2km tetrads during the breeding season (April to July) and winter period (November to February) was recorded following methods used by the BTO as part of their atlas surveys (BTO 2026, Balmer et al. 2013, Gillings *et al*. 2019). In Berkshire, data were collected from 394 tetrads between 2007 and 2011. In Cornwall, data were collected from 1050 tetrads between 2000 and 2009. In Devon, data were collected from 1858 tetrads between 2007 and 2013. In Hertfordshire, data were collected from 491 tetrads between 2007 and 2012. Across all four counties, there were a total of 3793 tetrads.

**Figure 1.**
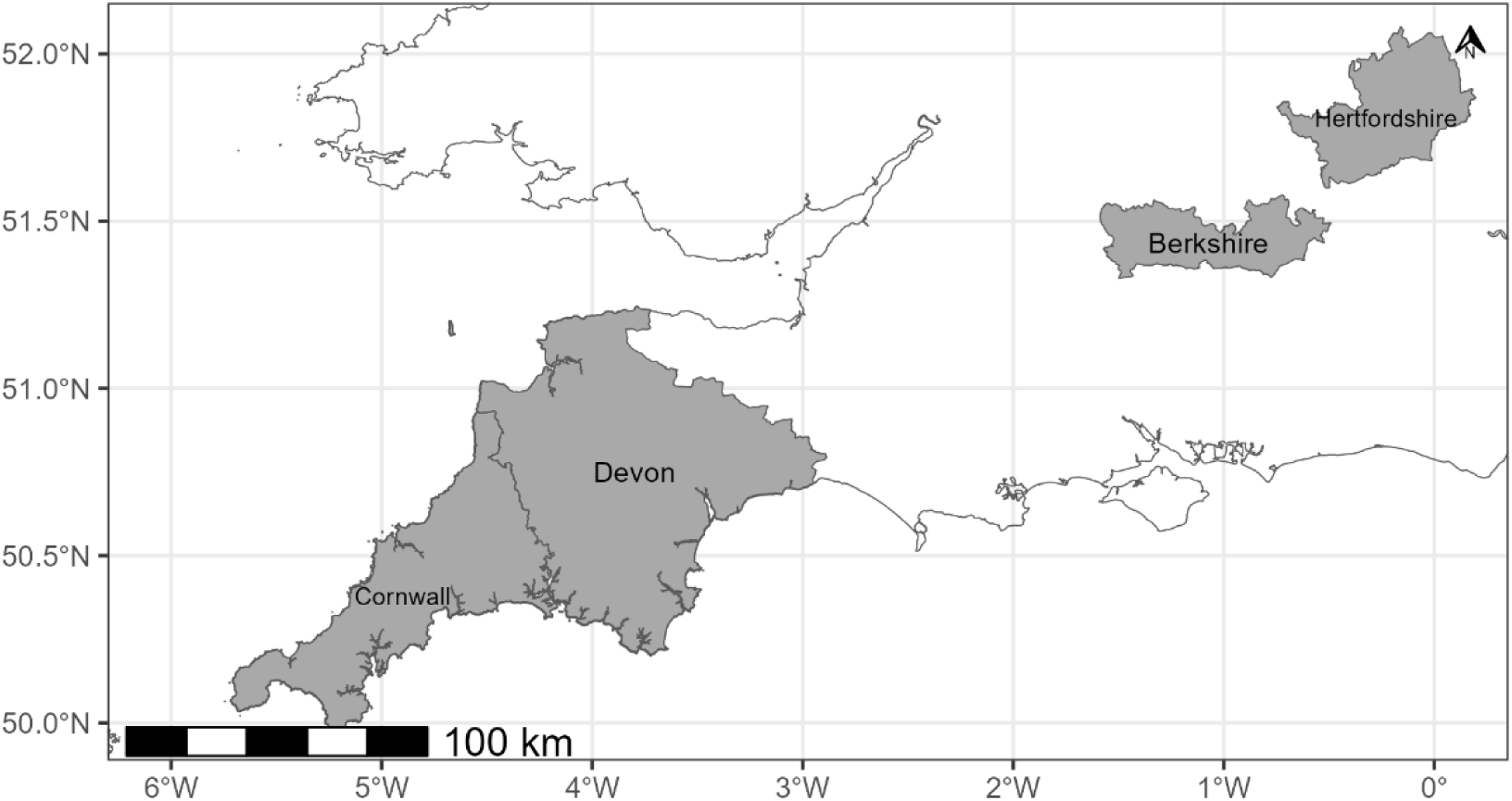
Map showing the location of each UK county for which the relative abundance of common pheasants was analysed. UK outline plotted using data from Countries (December 2021) Boundaries UK BUC, county outlines plotted using Counties and Unitary Authorities (May 2023) Boundaries UK BGC.

These data provide an abundance estimate for pheasants during the winter and breeding season in each tetrad, allowing for comparison between these two time periods. Between winter and the breeding season, the number of pheasants will have decreased overall due to shooting and natural mortality to approximately 15% of those released (Madden, Hall, and Whiteside, 2018). Therefore, we are interested in changes in the abundance of pheasants relative to the population within a county at a given time. Investigating this allows us to account for changes in population size due to mortality and releases.

### ii) Standardising abundance measures across counties

Since abundance was reported differently in each county, we standardised abundance measures across counties to get consistent, continuous measures across counties (Figure 2). Abundance data were provided as continuous values in Cornwall and remained unaltered. For Devon, Berkshire and Hertfordshire, reported abundance data were categorical with each category representing a range of abundance values. These categories were different for each of the three counties. To facilitate comparison among county data, we condensed these categories to a single midpoint value meaning each tetrad in each of these three counties had a single continuous value of abundance for winter and summer. Although this method allows us to analyse data from all four counties systematically and consistently, we acknowledge that using categorical measures in this way could be considered coarse and underestimate true uncertainty in parameter estimates.

**Figure 2.**
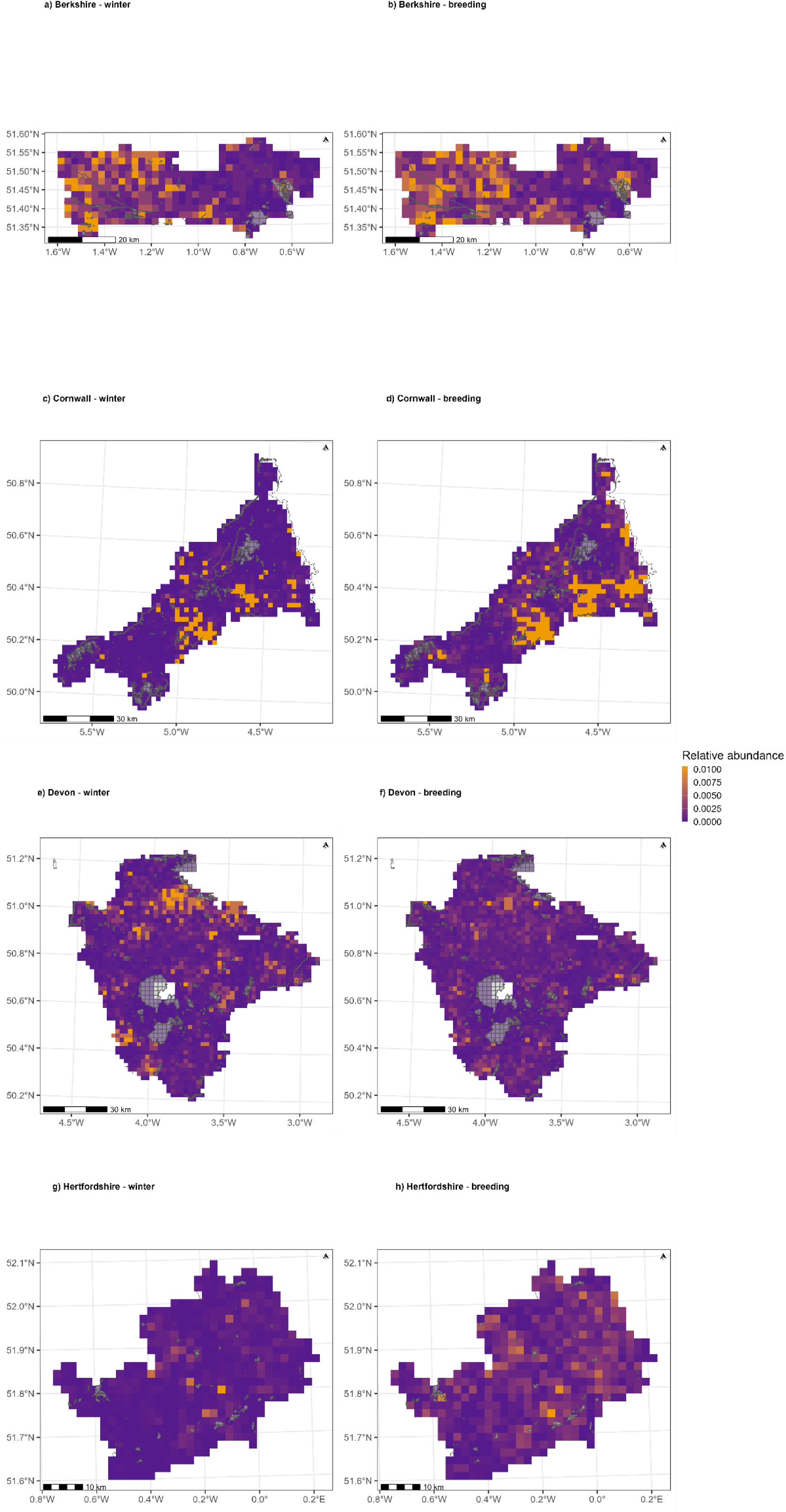
The relative pheasant abundance in each 2km^2^ tetrad in (a & b) Berkshire, (c & d) Cornwall, (e & f) Devon and (g & h) Hertfordshire. Relative abundances in winter (left column; a, c, e, g) and in the pheasant breeding season (right column; b, d, f, h) are both shown. Protected areas of ecologically relevant habitat type are overlayed in grey. The county outlines were plotted using data from Counties and Unitary Authorities (May 2023) Boundaries UK BGC.

### iii) Creating a measure of relative abundance

We were interested in changes in abundance relative to the total sample of pheasants alive in each county in each season. This would account for mass releases in late summer and high levels of mortality post release. We calculated these relative abundance measures using the following equation:

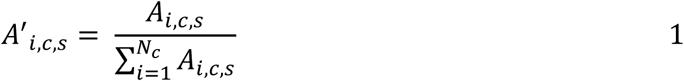

where:

- *A*_*i,c,s*_ and *A*′_*i,c,s*_: the absolute and relative abundance metric for the *i*th tetrad in county *c* during season *s*, respectively.
- *N*_*c*_: the total number of tetrads in county *c*.

This relative abundance metric can be interpreted as the proportion of the total population of pheasants in a single county in a single season that reside in a given tetrad. This metric would be constant for a given tetrad between time periods if there is no net immigration or emigration between tetrads, and survival rates are constant across the county.

If pheasants preferentially disperse to and reside in protected areas in a given season, then we would expect evidence of a positive relationship between this relative abundance metric and protected area coverage in that season, i.e. a higher proportion of a county’s population of pheasants can be found within protected areas than outside of protected areas. A finding of no relationship between relative abundance and protected area coverage suggests the same proportion of a county’s population can be found within protected areas as outside. A negative relationship suggests a smaller proportion of a county’s pheasant population can be found inside protected areas than outside. If we find a more positive relationship between relative abundance and protected areas coverage in the breeding season relative to the winter season (even if the overall relationship is still negative in both seasons), this may suggest that pheasants are preferentially dispersing into protected areas from the sites at which they are released.

### iv) Protected areas

Across the four counties examined, we calculated the proportion of each tetrad covered by protected areas (information on which was downloaded from the Government Digital Service’s dataset archive; https://ckan.publishing.service.gov.uk/dataset) that were ecologically relevant to pheasants. We used the 2007 United Kingdom Centre for Ecology and Hydrology (UKCEH) land cover map (25m x 25m spatial resolution; Morton *et al*., 2014) to determine the dominant habitat type in each protected area. Protected areas were classified as ecologically irrelevant to pheasants and removed from subsequent analysis if the dominant habitat type was one of the following: ocean, saltwater, freshwater, saltmarsh, bog, urban, suburban, inland rock, supra-littoral sediment, supra-littoral rock, littoral sediment or littoral rock. This meant protected areas mostly taken up by the following habitat types were retained: broadleaf woodland, coniferous woodland, arable/horticulture, fen/marsh/swamp, heather, heather grassland, acid grassland, rough grassland, improved grassland, neutral grassland and calcareous grassland. These remaining habitats are where pheasants are most commonly found in breeding bird surveys (Heywood *et al*., 2023). Occasionally a single designated site was composed of multiple distinct and disconnected polygons. In these instances, each polygon was treated as an independent protected area.

### v) Habitat data

Pheasants show distinct preferences for particular habitat types (e.g. Hill & Ridley 1987, Robertson et al. 1993a,b, Lachlan & Bray 1976). Therefore, we controlled for this by considering habitat coverage in each tetrad. Using the UKCEH land cover map (25m x 25m spatial resolution), we calculated the proportion of each tetrad covered by each of ten aggregate habitat classes: broadleaf woodland, coniferous woodland, arable, improved grassland, semi-natural grassland, mountain heath & bog, saltwater, freshwater, coastal and built-up areas and gardens (Morton *et* al., 2014), yielding ten continuous values between zero and one for each tetrad. We also calculated the Shannon diversity index for habitat in each tetrad, calculated as 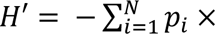 ln (*p*_*i*_), where *p*_*i*_is the proportion of each tetrad taken up by the *i*th habitat type, and *N* is the total number of possible habitat types (Ortiz-Burgos, 2016). Higher values of *H*′ indicate more habitat types in a tetrad.

### v) Statistical analyses

To analyse the relationship between protected area coverage and the relative abundance of pheasants, we used an occupancy-abundance model in a Bayesian framework. These models are a mixture of 1) occupancy models, which simultaneously estimate the probability that a target species occupies a given area (tetrad, in this case) and a probability that the target species is detected if present (Schaub & Kéry, 2022), and 2) abundance models, which estimate the abundance of the target species in occupied areas (here modelled using a Beta distribution; Bailey *et al*., 2014; Holt *et al*., 2002; Potts & Elith, 2006). The model formulation is:

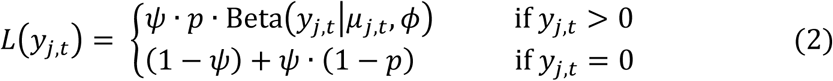

where *L*(*y*_*j,t*_) is the likelihood of observing the abundance metric (*y*) recorded in the *j*th tetrad in the *t*th season. If pheasants were observed in the *j*th tetrad in season *t*, the likelihood is the product of the probability a tetrad is occupied (ψ), the probability pheasants are detected in a tetrad (*p*), and the probability of observing the pheasant abundance metric under a beta distribution with mean (μ_*j,t*_) and precision (φ). If pheasants were not observed in a tetrad in a season, the likelihood is the probability a tetrad is not occupied (1 − ψ) added to the probability a tetrad is occupied (ψ) but the pheasants were not detected (1 − *p*). The probability of occupancy (ψ) and the probability of detection (*p*) are both assumed to vary between tetrads, and are each pulled from a normal distribution with a population mean and standard deviation estimated by the model. A full breakdown of the model can be found in the supplementary materials.

These models also estimate the effects that covariates have on pheasant abundance to determine which features of a tetrad causes pheasant abundance to increase or decrease. The covariates included in the model were county (Berkshire, Cornwall, Devon, Hertfordshire), time-period (winter/breeding season), the proportion of a tetrad covered by each of the ten habitat types (one fixed effect for each habitat type), the Shannon habitat diversity index, and the proportion of each tetrad covered by ecologically relevant protected areas. We included a two-way interaction term between time-period and protected area coverage to determine if the effect of protected area coverage differed between the winter period and breeding season. We also included a three-way interaction between county, season and protected area coverage to account for the possibility that the effect of protected area coverage, and its interaction with time, may be different for each county. A model was also run without these three-way interactions, to determine whether results changed in a model with fewer parameters. We also included an offset of the number of tetrads per county to account for any differences that county size would have on the relative abundance metric modelled.

All models were written in Stan (Carpenter *et al*., 2017) and compiled using Cmdstan (Stan Development Team, 2018) and the *cmdstanr* package (version 0.8.0; Gabry & Johnson, 2023) in R (version 4.4.0; R Core Team, 2022) using RStudio (version 2024.09.0; RStudio Team, 2020) and were run on Crop Diversity High Performance Cluster, described by Percival-Alwyn et al. (2024). Weakly informative priors were used for all parameters, and model convergence was checked by inspecting trace plots and ensuring that the potential scale reduction factor (*R̂*) ≈ 1. The 2.5% and 97.5% highest density limits (HDL) of posterior distributions are reported in square brackets throughout (e.g. [-0.01, 0.01]; Kruske, 2014; McElreath, 2020). Residuals were plotted spatially to inspect whether prediction accuracy was uniform across all counties and explore spatial autocorrelation. These residual plots, as well the priors used, and the full model output can be found in Table 3 in the Supplementary Materials.

## RESULTS

### i) Habitat type

The strongest effect on pheasant abundance was caused by the 10 habitat types (Table 1). There were positive relationships between relative abundance and the proportion of a tetrad occupied by broadleaf woodland (*β* = [0.00, 0.06], Figure 3), and arable (*β* = [0.00, 0.09]). The proportion of each tetrad occupied by built-up areas and gardens had a negative relationship with relative abundance (*β* = [-0.08, 0.00]).

**Table 1.** The lower and upper highest density limits (HDL) of the posterior distribution of the effect size of the proportion of each tetrad occupied by all ten land-use habitat types, as well as the effect of the Shannon habitat diversity index. Bold text indicates effect sizes where the HDL does not overlap zero.

| *Habitat type* | *2.5% HDL of effect size* | *97.5% HDL of effect size* |
| --- | --- | --- |
| **Broadleaf woodland** | **0.00** | **0.06** |
| Coniferous woodland | -0.03 | 0.02 |
| **Arable** | **0.00** | **0.09** |
| Improved grassland | -0.04 | 0.05 |
| Semi-natural grassland | -0.06 | 0.03 |
| Mountain, heath and bog | -0.08 | 0.04 |
| Saltwater | -0.03 | 0.03 |
| Freshwater | -0.04 | 0.01 |
| Coastal | -0.03 | 0.03 |
| **Built-up areas and gardens** | **-0.08** | **0.00** |
| Shannon habitat diversity index | -0.04 | 0.03 |

**Figure 3.**
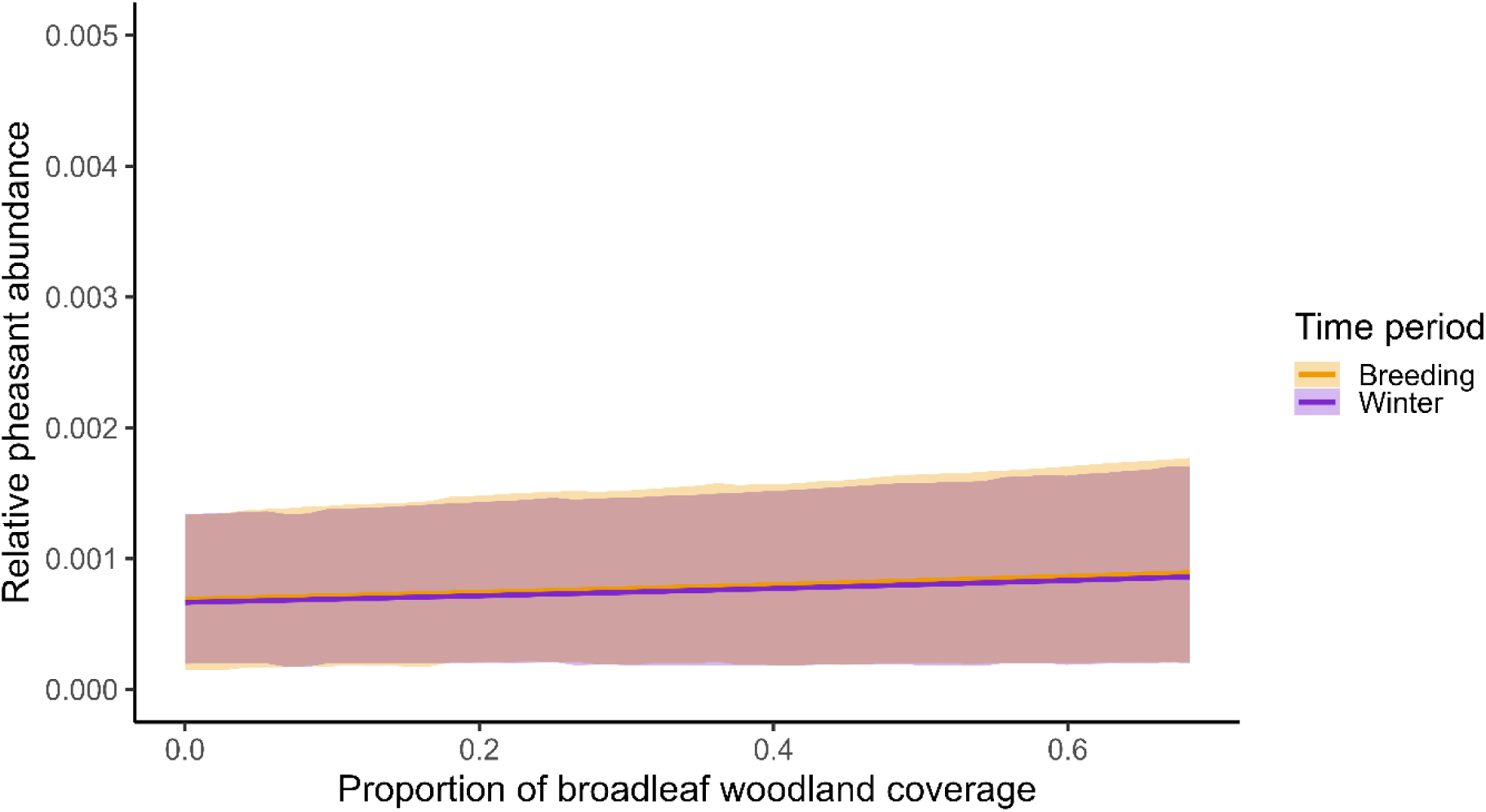
The model predicted relationship between the proportion of broadleaf woodland coverage and mean relative pheasant abundance (coloured lines), while other model predictors are held constant at their mean value. The shaded areas denote the 95% highest density interval around the predicted mean for each time period.

### ii) Protected area coverage

There was no evidence of an effect of the proportion of each tetrad occupied by protected areas on pheasant abundance (*β* = [-0.06, 0.02], Figure 4) with approximately equal abundances recorded in tetrads containing high and low proportions of protected area coverage. There was no evidence of a two-way interaction between protected area coverage and time period (*β* = [-0.09, 0.08]), suggesting the (lack of) effect of protected area coverage is consistent between winter and the breeding season. There was also no evidence of a three-way interaction between protected area coverage, time period, and county (Table 2), suggesting that the distribution patterns are consistent across the four counties that we considered. The model that did not include the three-way interactions found the same results as the model that included them (Table 4, ESM). The full results from the occupancy-abundance model can be found in Table 3 in the supplementary materials.

**Figure 4.**
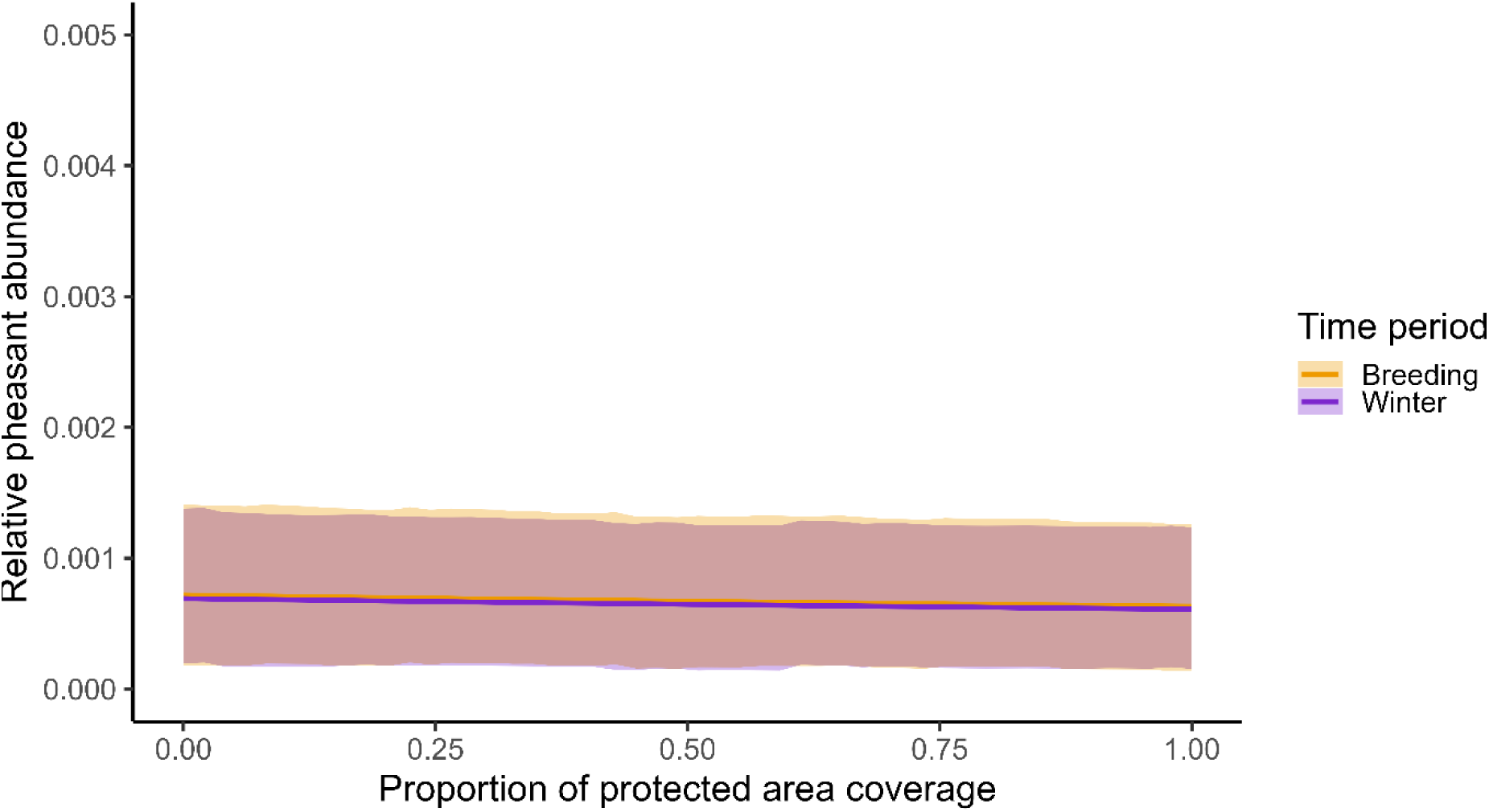
The model predicted relationship between protected area coverage and mean relative abundance is shown (coloured lines), while other model predictors are held constant at their mean value. The shaded areas denote the 95% highest density interval around the predicted mean for each time period.

**Table 2.** The highest density limits (HDL) of the posterior distribution of the effect size of the proportion of each tetrad occupied by suitable protected areas, the time period in which the abundance survey took place, as well as the two- and three-way interactions between protected area coverage, time period and county.

| *Fixed effect* | *2.5% HDL of effect size* | *97.5% HDL of effect size* |
| --- | --- | --- |
| Protected area (PA) coverage | -0.06 | 0.02 |
| Time period: Winter → Breeding | -0.01 | 0.04 |
| PA coverage × Time period: Winter → Breeding | -0.09 | 0.08 |
| PA coverage × Time period: Winter × Berkshire | -0.18 | 0.15 |
| PA coverage × Time period: Winter × Cornwall | -0.14 | 0.14 |
| PA coverage × Time period: Winter × Devon | -0.15 | 0.14 |
| PA coverage × Time period: Winter × Hertfordshire | -0.14 | 0.15 |
| PA coverage × Time period: Breeding × Berkshire | -0.17 | 0.17 |
| PA coverage × Time period: Breeding × Cornwall | -0.14 | 0.15 |
| PA coverage × Time period: Breeding × Devon | -0.15 | 0.15 |
| PA coverage × Time period: Breeding × Hertfordshire | -0.15 | 0.14 |

## DISCUSSION

We used relative pheasant abundance measures calculated from tetrad survey data to determine if there was a relationship between pheasant abundance and ecologically suitable protected area coverage. We also investigated whether any relationship was consistent between seasons and across counties. We did this to determine if there is any evidence suggesting pheasants preferentially disperse into and reside in protected areas, especially during the spring/summer once management actions associated with the shooting season have ceased or declined. This is of particular interest if pheasants disperse into protected areas and cause negative ecological effects in these sensitive areas.

The relationship between broadleaf woodland, arable and built-up area coverages and relative pheasant abundance reflect known habitat preferences of pheasants in the UK (e.g. Hill & Ridley 1987, Robertson et al. 1993a,b, Lachlan & Bray 1976). The positive relationship between relative pheasant abundance and broadleaf woodland coverage and arable coverage is expected given these habitats are where pheasants are most often released (Robertson *et al*., 2017; Heywood *et al*., 2023), and it is their preferred habitat (Ashoori *et al*., 2018). These results demonstrate that our broad-scale approach has been able to capture known ecological drivers of pheasant abundance by detecting preferences for land cover type.

We found no evidence of a relationship between protected area coverage and relative pheasant abundance. This lack of a relationship was consistent across all four counties and time periods. This suggests that pheasants do not preferentially disperse into or reside in protected areas and disperse equally into protected and non-protected areas. This may be because protected areas do not provide habitat that is more preferable than surrounding unprotected areas for pheasants, such as farmland or managed woodland. Alternatively, or additionally, protected areas may have high levels of human recreational disturbance from visitors to naturally beautiful or interesting sites (Mammides, 2020), making them less attractive to dispersing gamebirds, despite the fact that hunting within them is reduced.

We may also observe a lack of preferential movement into protected areas as pheasants are unlikely to be released uniformly across a county with respect to protected areas. Since 1981, consent has been required to release or manage gamebirds (including supplementary feeding or siting release pens) on SSSIs. Since 2021 (after the data used in the study were collected), releases in 500m buffer zones around SACs and SPAs can only be conducted under general or individual licences. The requirements of additional licence or consenting requirements are believed to dissuade game managers from releasing in such areas (BASC, 2025). Therefore, game managers might now be more likely to release outside protected area boundaries, or outside the buffer zones surrounding protected areas, or they may release them in these areas at lower densities than elsewhere so as to comply with general licence requirements. It is therefore possible that tetrads with high protected area coverage also have lower densities of pheasant release occurring within them, and this may also be true of when the data used in this study were collected. One may expect that this would be detected in our analysis, with pheasant abundance being higher in tetrads with low protected area coverage. However, without more precise estimates of the number and location of release sites in a county, it is difficult to determine how much of this signal we would expect to be able to detect.

Pheasants are generally unlikely to move large distances away from the site of their release, either due to a naturally sedentary nature or more likely in the UK, because of deliberate management to retain them close to release sites so that they are available to hunt (Sage *et al*. 2021). Therefore, even if pheasants would preferentially reside in protected areas, they may not encounter this opportunity due to restrictions on release sites and limited movement. To investigate this phenomenon introduced by release restrictions and pheasant dispersal, further work needs to be carried out using information on the number and density of pheasant releases occurring in each tetrad. Currently, such accurate and precise data on releases are lacking.

Due to the spatial scale of the data used, we can only investigate pheasant dispersal between tetrads. Because we focus on a 2km x 2km tetrad as our spatial unit, we would be unable to detect dispersal occurring if most dispersing birds remain within the same tetrad into which they were released, and data collected at a finer spatial resolution would be needed to investigate this. Most pheasants are unlikely to move long distances from their site of release (Sage *et al*. 2021) and so there may be some dispersal not detected by this analysis. However, we expect release sites to be randomly distributed within each tetrad in our analysis, meaning we are still likely to detect pheasant dispersal into adjacent tetrads. Further work should seek to carry out similar analyses at a finer spatial scale, while accounting for the location of known release sites.

The results of our investigation suggest that pheasants are not preferentially dispersing into or residing in protected areas at any time of year. Therefore, there is no evidence to suggest that any ecological impact wrought by pheasants is likely to happen more in protected areas than non-protected areas. However, it should be noted that our findings do not suggest that direct negative ecological impacts caused by released pheasants (Madden & Sage 2020, Mason et al. 2020, Sage et al. 2020, Madden 2023) can not occur in protected areas. The numbers and densities of pheasants detected in protected areas, even when equal to rather than exceeding, those in surrounding unprotected areas may still lead to negative ecological impacts. Indeed, a similar number or density of birds may cause greater ecological change in especially important or sensitive protected areas or on the particular features contained in them. However, the nature and relative scale of ecological change wrought by released gamebirds within and outside protected areas is unknown, particularly when considering the impact of birds at low density (Madden *et al*. 2026). Future research should investigate whether these patterns hold at a finer spatial scale using information from known releases, as well as explore the implications of gamebird management practices for both pheasant populations and biodiversity within protected areas.

## Supporting information

ESM

## ACKNOWLEDGEMENTS

This work was made possible due to funding from the Animal and Plant Health Agency (APHA) and the British Association for Shooting and Conservation (BASC). The Game and Wildlife Conservation Trust (GWCT) provided knowledge and feedback to the project. Thanks to Marnie Lovejoy & Cat McNicol (BASC), Graham Smith, Rufus Sage, Iain McKendrick and Stephen Catterall for feedback on drafts of the manuscript. The authors acknowledge Research Computing at the James Hutton Institute for providing computational resources and technical support for the “UK’s Crop Diversity Bioinformatics HPC” (BBSRC grants BB/S019669/1 and BB/X019683/1), use of which has contributed to the results reported within this paper. We are very grateful to the volunteer bird recorders and atlas compilers who produced the county bird atlas’s.

## AUTHOR CONTRIBUTIONS

J.R.M. and L.O-H. conceived of the presented idea. J.R.M. developed the theory and supervised the project, L.O-H developed the data cleaning pipeline, and J.A.W. developed the analysis pipeline, and data/results presentation. J.A.W. and J.R.M. wrote the final manuscript. All authors discussed and contributed to the final manuscript.

## DATA AVAILABILITY

The data supporting this MS and the R code used in the analyses are available from https://github.com/JoeAWilde/BTO_Tetrad_Analysis. They will be placed in a permanent repository on acceptance of the paper.

