## Supplementary material for "A spatial analysis of Common Pheasant (*Phasianus colchicus*) abundance with reference to Protected Area coverage in England": ESM

| | **Posterior values** | | **95% Highest Density Limits** | | | **Effective sample size** | | |
|---|---|---|---|---|---|---|---|---|
| **Parameter** | **Mean** | **SD** | **Lower** | **Upper** | **R-hat** | **Bulk** | **Tail** | **Prior distribution** |
| Pr(detection) | 0.97 | 0.01 | 0.94 | 0.99 | 1 | 368 | 871 | Normal(0.5, 1) |
| **Pr(occupied) (logit)** | | | | | | | | |
| Population mean | 2.91 | 0.32 | 2.36 | 3.57 | 1 | 410 | 891 | Normal(0, 1) |
| Population SD | 2.70 | 0.27 | 2.21 | 3.25 | 1 | 465 | 1052 | Exponential(1) |
| **Abundance (logit)** | | | | | | | | |
| **Intercept** | | | | | | | | |
| Population mean | -7.37 | 0.47 | -8.28 | -6.46 | 1 | 2398 | 2504 | Normal(0, 1) |
| Population SD | 0.01 | 0.01 | 0.00 | 0.03 | 1 | 3924 | 1876 | Exponential(1) |
| **Covariates** | | | | | | | | |
| County: Berkshire → Cornwall | -0.02 | 0.03 | -0.09 | 0.04 | 1 | 2506 | 2358 | Normal(0, 0.1) |
| County: Berkshire → Devon | 0.03 | 0.05 | -0.07 | 0.13 | 1 | 2286 | 2681 | Normal(0, 0.1) |

| | | | | | | | | |
|---|---|---|---|---|---|---|---|---|
| County: Berkshire → Hertfordshire | -0.02 | 0.02 | -0.05 | 0.02 | 1 | 5634 | 3223 | Normal(0, 0.1) |
| Time period: winter → breeding | 0.02 | 0.01 | -0.01 | 0.04 | 1 | 10437 | 2822 | Normal(0, 0.1) |
| Habitat: Broadleaf woodland | 0.03 | 0.02 | 0.00 | 0.06 | 1 | 3741 | 3236 | Normal(0, 0.1) |
| Habitat: Coniferous woodland | 0.00 | 0.01 | -0.03 | 0.02 | 1 | 6141 | 2864 | Normal(0, 0.1) |
| Habitat: Arable | 0.04 | 0.02 | 0.00 | 0.09 | 1 | 2702 | 3120 | Normal(0, 0.1) |
| Habitat: Improved grassland | 0.00 | 0.02 | -0.04 | 0.05 | 1 | 3308 | 3298 | Normal(0, 0.1) |
| Habitat: Semi-natural grassland | -0.01 | 0.02 | -0.06 | 0.03 | 1 | 4335 | 3283 | Normal(0, 0.1) |
| Habitat: Mountain, heath and bog | -0.01 | 0.02 | -0.06 | 0.04 | 1 | 5354 | 3073 | Normal(0, 0.1) |
| Habitat: Saltwater | 0.00 | 0.02 | -0.03 | 0.03 | 1 | 8660 | 2974 | Normal(0, 0.1) |
| Habitat: Freshwater | -0.02 | 0.01 | -0.04 | 0.01 | 1 | 7184 | 3311 | Normal(0, 0.1) |

| | | | | | | | | |
|---|---|---|---|---|---|---|---|---|
| Habitat: Coastal | 0.00 | 0.02 | -0.03 | 0.03 | 1 | 5327 | 3244 | Normal(0, 0.1) |
| Habitat: Built-up areas and gardens | -0.04 | 0.02 | -0.08 | 0.00 | 1 | 3129 | 3163 | Normal(0, 0.1) |
| Shannon habitat diversity index | 0.00 | 0.02 | -0.04 | 0.03 | 1 | 3652 | 3179 | Normal(0, 0.1) |
| Protected area (PA) coverage | -0.02 | 0.02 | -0.06 | 0.02 | 1 | 6073 | 3273 | Normal(0, 0.1) |
| **Two-way interactions** | | | | | | | | |
| PA coverage × Time period: winter | 0.00 | 0.04 | -0.09 | 0.08 | 1 | 3781 | 2941 | Normal(0, 0.1) |
| **Three-way interactions** | | | | | | | | |
| PA coverage × Time period: winter × County: Berkshire | 0.00 | 0.09 | -0.18 | 0.15 | 1 | 5910 | 2663 | Normal(0, 0.1) |
| PA coverage × Time period: winter × County: Cornwall | 0.00 | 0.07 | -0.14 | 0.14 | 1 | 4589 | 2881 | Normal(0, 0.1) |
| PA coverage × Time period: winter × County: Devon | 0.00 | 0.08 | -0.15 | 0.14 | 1 | 4461 | 2955 | Normal(0, 0.1) |
| PA coverage × Time period: winter × County: Hertfordshire | 0.00 | 0.07 | -0.14 | 0.15 | 1 | 5297 | 3010 | Normal(0, 0.1) |

| | | | | | | | | |
|---|---|---|---|---|---|---|---|---|
| PA coverage × Time period: breeding × County: Berkshire | 0.00 | 0.09 | -0.17 | 0.17 | 1 | 5238 | 3218 | Normal(0, 0.1) |
| PA coverage × Time period: breeding × County: Cornwall | 0.00 | 0.07 | -0.14 | 0.15 | 1 | 5246 | 3111 | Normal(0, 0.1) |
| PA coverage × Time period: breeding × County: Devon | 0.00 | 0.08 | -0.15 | 0.15 | 1 | 4774 | 2803 | Normal(0, 0.1) |
| PA coverage × Time period: breeding × County: Hertfordshire | 0.00 | 0.07 | -0.15 | 0.14 | 1 | 5523 | 2803 | Normal(0, 0.1) |
| **Per-county offset** | | | | | | | | |
| log(No. tetrads) | -0.51 | 0.07 | -0.63 | -0.38 | 1 | 2405 | 2322 | Normal(0, 0.1) |
| **Family-specific parameters** | | | | | | | | |
| Phi (precision) | 11511 | 373 | 10803 | 12272 | 1 | 10761 | 2691 | Exponential(0.1) |

Table 3 – model output for the full occupancy-abundance model, as well as the prior distribution used for each parameter.

a)

c)

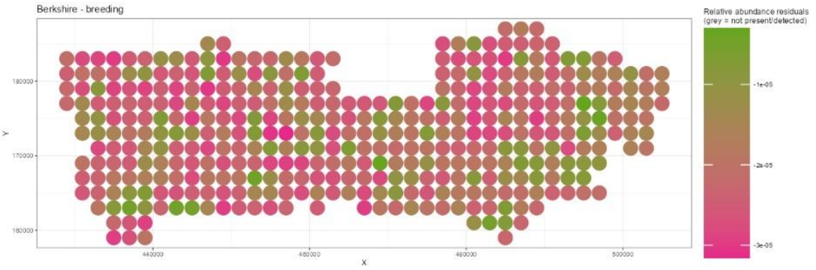
Berkshire - breeding
Relative abundance residuals
(grey = not present/detected)
-1e-05
-2e-05
-3e-05
180000
170000
160000
Y
440000
460000
480000
500000
X

b)

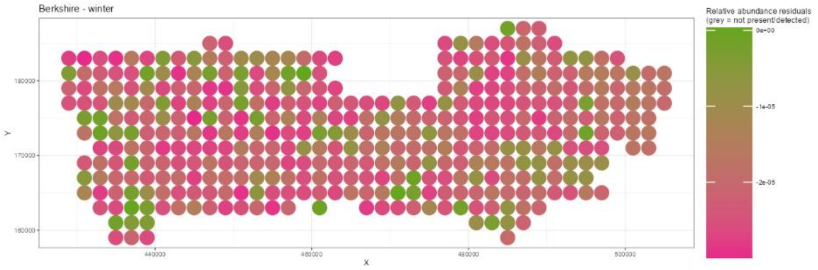
Berkshire - winter
Relative abundance residuals
(grey = not present/detected)
0e+00
-1e-05
-2e-05
180000
170000
160000
Y
440000
460000
480000
500000
X

d)

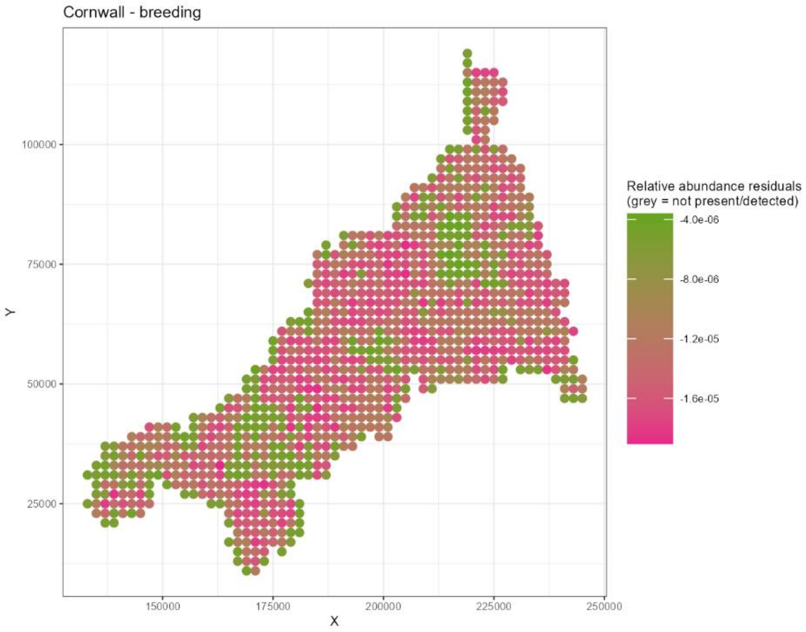
Cornwall - breeding
Relative abundance residuals
(grey = not present/detected)
-4.0e-06
-8.0e-06
-1.2e-05
-1.6e-05
100000
75000
50000
25000
Y
150000
175000
200000
225000
250000
X

e)

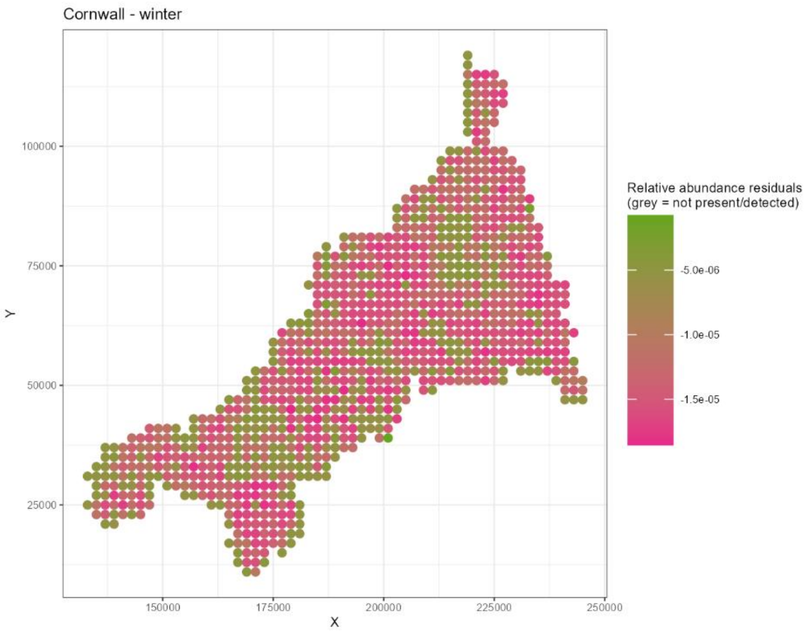

Cornwall - winter
100000
75000
50000
25000
Y
150000
175000
200000
225000
250000
X
Relative abundance residuals
(grey = not present/detected)
-5.0e-06
-1.0e-05
-1.5e-05


f)

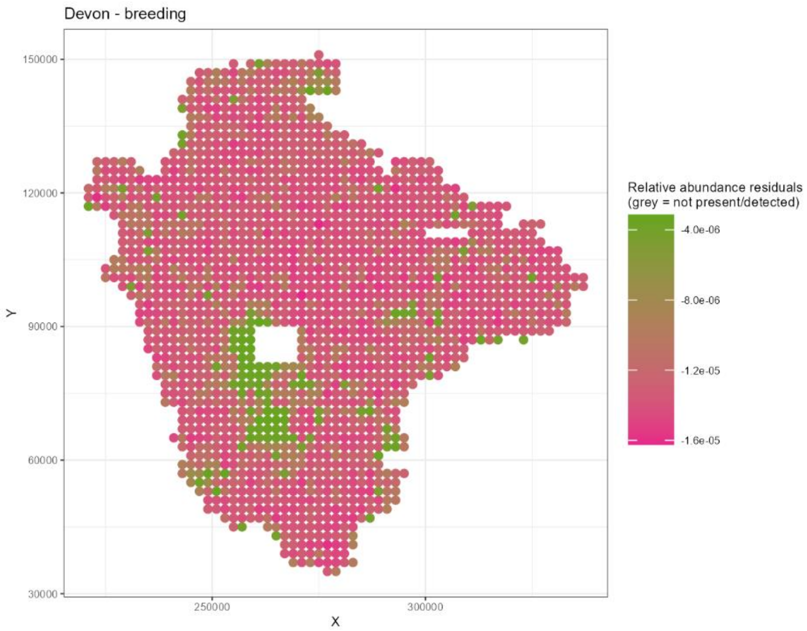

Devon - breeding
150000
120000
90000
60000
30000
Y
250000
300000
X
Relative abundance residuals
(grey = not present/detected)
-4.0e-06
-8.0e-06
-1.2e-05
-1.6e-05

g)

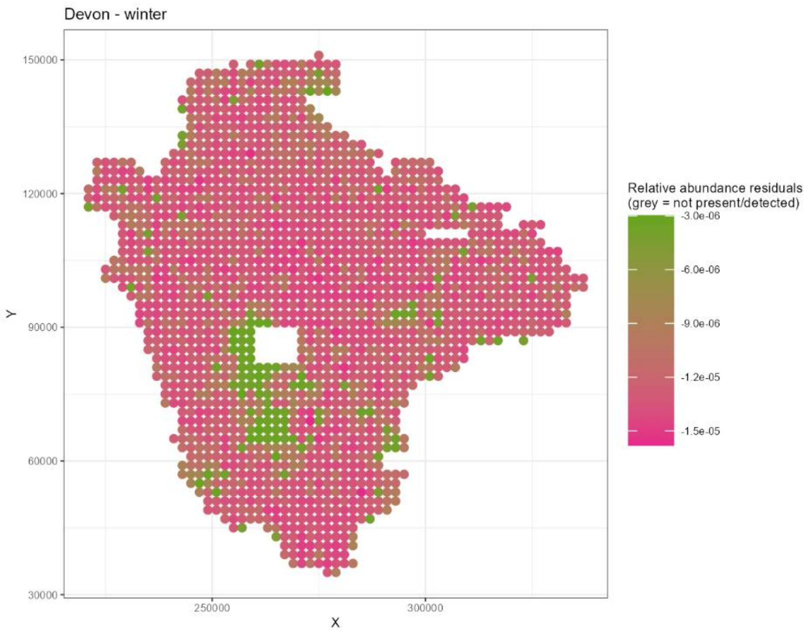
Devon - winter
150000
120000
90000
60000
30000
Y
250000
300000
X
Relative abundance residuals
(grey = not present/detected)
-3.0e-06
-6.0e-06
-9.0e-06
-1.2e-05
-1.5e-05

h)

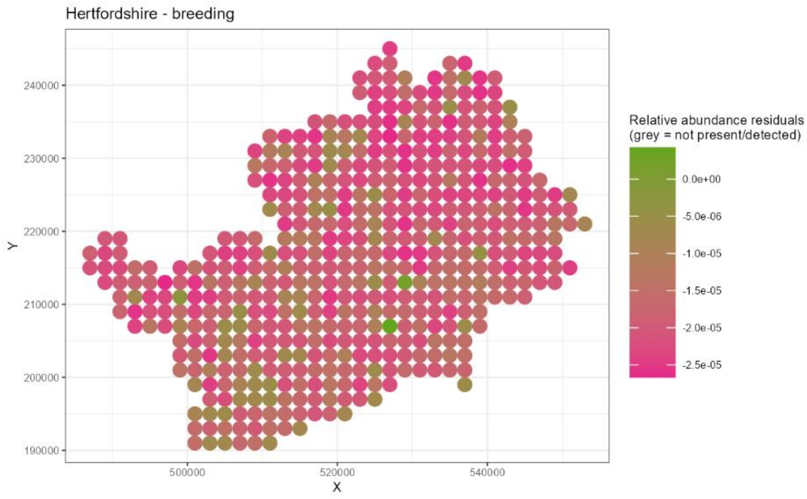
Hertfordshire - breeding
240000
230000
220000
210000
200000
190000
Y
500000
520000
540000
X
Relative abundance residuals
(grey = not present/detected)
0.0e+00
-5.0e-06
-1.0e-05
-1.5e-05
-2.0e-05
-2.5e-05

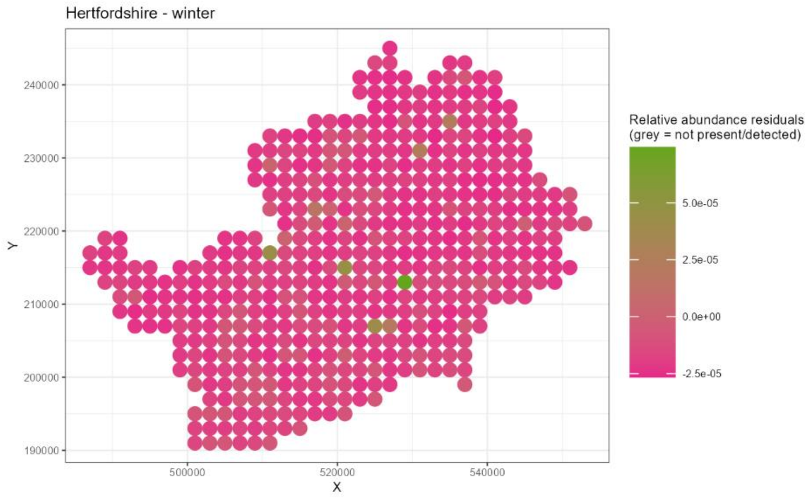
Hertfordshire - winter
240000
230000
220000
210000
200000
190000
Y
500000
520000
540000
X
Relative abundance residuals
(grey = not present/detected)
5.0e-05
2.5e-05
0.0e+00
-2.5e-05

**Figure 5** – the difference between the mean of the model predicted relative abundance and the true relative abundance measure for each tetrad (residuals), plotted spatially across Berkshire (a – breeding season, b – winter), Cornwall (c – breeding season, d – winter), Devon (e – breeding season, f – winter), and Hertfordshire (g – breeding season, h – winter).

| | **Posterior values** | | **95% Highest Density Limits** | | | **Effective sample size** | |
|---|---|---|---|---|---|---|---|
| **Parameter** | **Mean** | **SD** | **Lower** | **Upper** | **R-hat** | **Bulk** | **Tail** |
| Pr(detection) | 0.97 | 0.01 | 0.94 | 0.99 | 1 | 276 | 839 |
| **Pr(occupied) (logit)** | | | | | | | |
| Population mean | 2.92 | 0.37 | 2.33 | 3.66 | 1 | 325 | 1029 |
| Population SD | 2.72 | 0.30 | 2.21 | 3.34 | 1 | 433 | 1160 |
| **Abundance (logit)** | | | | | | | |
| **Intercept** | | | | | | | |
| Population mean | -7.36 | 0.47 | -8.31 | -6.46 | 1 | 4114 | 4506 |
| Population SD | 0.01 | 0.01 | 0.00 | 0.03 | 1 | 5700 | 3316 |
| **Covariates** | | | | | | | |
| County: Berkshire → Cornwall | -0.02 | 0.03 | -0.09 | 0.04 | 1 | 4069 | 4999 |
| County: Berkshire → Devon | 0.03 | 0.05 | -0.07 | 0.13 | 1 | 3761 | 4342 |
| County: Berkshire → Hertfordshire | -0.02 | 0.02 | -0.05 | 0.02 | 1 | 10153 | 4758 |

| | | | | | | | |
|---|---|---|---|---|---|---|---|
| Time period: winter → breeding | 0.02 | 0.01 | -0.01 | 0.04 | 1 | 16225 | 4093 |
| Habitat: Broadleaf woodland | 0.03 | 0.02 | 0.00 | 0.06 | 1 | 6256 | 5557 |
| Habitat: Coniferous woodland | 0.00 | 0.01 | -0.03 | 0.02 | 1 | 9318 | 4706 |
| Habitat: Arable | 0.04 | 0.02 | 0.00 | 0.08 | 1 | 4480 | 5072 |
| Habitat: Improved grassland | 0.00 | 0.02 | -0.04 | 0.05 | 1 | 4906 | 4839 |
| Habitat: Semi-natural grassland | -0.01 | 0.02 | -0.06 | 0.03 | 1 | 7033 | 4761 |
| Habitat: Mountain, heath and bog | -0.01 | 0.02 | -0.06 | 0.04 | 1 | 8820 | 4833 |
| Habitat: Saltwater | 0.00 | 0.02 | -0.03 | 0.03 | 1 | 13784 | 3962 |
| Habitat: Freshwater | -0.01 | 0.01 | -0.04 | 0.01 | 1 | 11513 | 4527 |
| Habitat: Coastal | 0.00 | 0.02 | -0.03 | 0.03 | 1 | 8149 | 4882 |
| Habitat: Built-up areas and gardens | -0.04 | 0.02 | -0.08 | 0.00 | 1 | 5124 | 5137 |
| Shannon habitat diversity index | 0.00 | 0.02 | -0.04 | 0.03 | 1 | 6992 | 4804 |
| Protected area (PA) coverage | -0.02 | 0.02 | -0.06 | 0.02 | 1 | 11581 | 4892 |
| **Two-way interactions** | | | | | | | |
| PA coverage × Time period: winter | 0.00 | 0.02 | -0.04 | 0.03 | 1 | 15774 | 4594 |
| **Per-county offset** | | | | | | | |
| log(No. tetrads) | -0.51 | 0.07 | -0.63 | -0.37 | 1 | 4124 | 4482 |
| **Family-specific parameters** | | | | | | | |
| Phi (precision) | 11509.32 | 351.33 | 10825.20 | 12203.10 | 1 | 16644 | 3795 |

Table 4 – model output for the occupancy-abundance model without three-way interactions between county, time period and protected area coverage. Also shown is the prior distribution used for each parameter.

**Model outline**

$$L(y_{j,t}) = \begin{cases} \psi \cdot p \cdot \mathrm{Beta}(y_{j,t} | \mu_{j,t}, \phi) & \text{if } y_{j,t} > 0 \\ (1-\psi) + \psi \cdot (1-p) & \text{if } y_{j,t} = 0 \end{cases}$$

$$\mathrm{logit}(\psi_j) = \alpha_j^{\mathrm{occ}}$$

$$\mathrm{logit}(p) = \delta$$

$$\mathrm{logit}(\mu_{j,t}) = \alpha_j^{\mathrm{abund}} + \sum_{k=1}^{K} \beta_k X_{j,t,k}$$

$$\alpha_j^{\mathrm{occ}} \sim \mathrm{Normal}(\upsilon^{\mathrm{occ}}, \sigma^{\mathrm{occ}}) \qquad (3)$$

$$\upsilon^{\mathrm{occ}} \sim \mathrm{Normal}(0, 1)$$

$$\sigma^{\mathrm{occ}} \sim \mathrm{Exponential}(1)$$

$$\delta \sim \mathrm{Normal}(0.5, 1)$$

$$\alpha_j^{\mathrm{abund}} \sim \mathrm{Normal}(\upsilon^{\mathrm{abund}}, \sigma^{\mathrm{abund}})$$

$$\upsilon^{\mathrm{occ}} \sim \mathrm{Normal}(0, 1)$$

$$\sigma^{\mathrm{occ}} \sim \mathrm{Exponential}(1)$$

$$\beta_k \sim \mathrm{Normal}(0, 0.1)$$

$$\phi \sim \mathrm{Exponential}(0.1)$$

where:

- $y_{j,t}$: the pheasant abundance metric for the $j$th tetrad for the $t$th season.
- $\psi_j$: the probability of occupancy for the $j$th tetrad.
- $p$: the probability of detecting pheasants in a tetrad.

- $\mu_{j,t}$ and $\phi$: the mean and overall precision (respectively) of a beta distribution modelling the pheasant abundance metric for the $j$th tetrad for the $t$th season.
- $K$: total number of predictors.
- $X_{j,t,k}$: the value of the $k$th predictor (log(no. of tetrads) included in predictors) for the $j$th tetrad for the $t$th season.
